# The role of directionality in determining spatiotemporal tau pathology differs between AD-Like and Non-AD-Like mouse models

**DOI:** 10.1101/2020.11.06.371625

**Authors:** Christopher Mezias, Ashish Raj

## Abstract

**Introduction:** Current research indicates divergent spatiotemporal tauopathy progression between conditions and implicates transsynaptic connectome-based spread as a main mechanism. We examine tauopathy and connectome interactions to investigate why different spatiotemporal patterns of pathology arise.

**Methods:** We test whether divergent spatiotemporal tau pathology patterns from 15 mouse-model datasets can be explained by a directional bias in tau transmission along fiber tracts via a mathematical model called Directed Network Transmission (DNT).

**Results:** Amyloid-comorbid tauopathic mouse models meant to mimic AD demonstrate spatiotemporal tauopathy patterns consistent with retrograde direction spread biases. Non-amyloid-comorbid mice demonstrate no consistent spread biases. Further, canonically ‘early’ tau pathology regions in AD are implicated as having earliest pathology in a simulation with random tauopathy seeding locations with retrograde biased spread.

**Discussion:** These results implicate directional biases in tau pathology spread along fiber tracts as a strong candidate explanation for divergent spatiotemporal tau progression between conditions.

## Introduction

Characterizing the spatiotemporal pattern of pathology development is a longstanding aim of degenerative disease research, beginning with Braak Staging of hallmarked protein pathology for Alzheimer’s Disease (AD) neurofibrillary tangles (NFT) (Braak & Braak, 1991). There are important differences in disease progression, however, as AD tau tangles are first detectable in the entorhinal cortex (EC) and locus coerules (LC; Braak & Braak, 1991), while other tauopathic conditions can exhibit early pathology in the orbitofrontal cortex (OFC), the amygdala (AMY), and parts of the basal forebrain such as the nucleus accumbens (NAcc) and caudate nuclues (CN), (Burrell, 2014; Chiba, 2013). The differences in spatiotemporal protein progression lead to differences in symptomatic progression, as the as AD patients experience deficits in spatial and working memory, long-term recall, and navigation (Braak & Tredici, 2011), while those with other tauopathic conditions can demonstrate increased anxiety and aggression (Burrell, et al., 2014). Much effort is accordingly going into characterizing, predicting, and explaining the divergent spatiotemporal development of tau pathology across different tauopathic conditions.

Attempts to explain the spatiotemporal development of tau pathology and volumetric loss observed in patients with different degenerative tauopathic conditions take three approaches. First, the cellular mechanisms of hyperphosphorylated tau deduced from vitro and in vivo research appear to be converging on intercellular transsynaptic spread as a likely candidate (Holmes, 2014; Tai, 2014, Ahmed, 2014; Clavaguera, 2009). Second, clinical studies confirm that the two disease hallmarks, regional tau pathology and volumetric loss severity, are correlated (Ossenkoppele, 2015), but that their spatiotemporal development patterns are different between conditions, as are the brain regions showing early disease vulnerability (Braak & Tredici, 2011; Burrell, 2014). Third, mathematical network transmission (NT) models of pathology spread along anatomical connectivity successfully recapitulate the spatiotemporal regional volumetric loss in patients’ brains. These models show great promise in predicting later disease patterns from early timepoints (Raj, 2012; Raj, 2015; Zhou, 2012).

Despite these advances, the questions of how tauopathies can progress in divergent manners in different conditions remains unaddressed, and two alternative hypotheses may be discerned: 1) that tau pathology seeds in specific and separate locations in different syndromes (the Selective Vulnerability hypothesis; SVb; Seeley, et al., 2009), and 2) that pathological tau protein transits with a different mechanism in a disorder-dependent manner, similar to the Molecular Nexopathy hypothesis, originally posited by Warren, et al., 2013. The SVb hypothesis originally held much sway in degenerative disease research, but recent studies call into question whether there are specific pathology initiation loci and have specifically undermined regions such as the LC or the EC (Iba, et.al., 2013; Arendt, et.al., 2015) as disease initiators. In the last two decades, however, accumulating evidence highlights important and possibly condition altering differences between pathological tau species between degenerative conditions. While AD has both 3R and 4R tau isoforms present in NFTs, other tauopathic conditions exhibit misfolded 4R or 3R isforms exclusively (Higuchi, Trojanowski, & Lee, 2002). Gross tangle structure is also different across diseases, with AD tau fibrils forming NFTs and pre-helical formations (PHFs), Pick’s Disease tau forming PBs, and PSP, CBD, and FTD oligomers forming more abundant NTs (Clavaguera, et al., 2013). At the molecular level, sites of tau hyperphosphorylation tend to be different across diseases (Hanger, Anderton, & Noble, 2009).

Given the clear differences between pathological tau protein species involved in each disorder, we posit a novel hypothesis: *that divergent spatiotemporal patterns of development between different tauopathic disorders may not be due to consistent and divergent starting locations (brain region SVb to tauopathic spread or initiation), but rather are due to biochemical protein differences that are responsible for changing the mechanism of transsynaptic transmission, and therefore changing the preferred direction of transcellular spread (MNx specifically manifesting in different relative to axon polarity spread biases among different pathological tau species).*

We accordingly undertake the current study to investigate whether either intrinsic regional SVb or a MNx-like mechanism positing transsynaptic spread direction biases due to differences between pathological tau species can offer compelling explanations for spatiotemporal tau progression patterns. We employ a mathematical Network Transmission (NT) model of pathology spread, expanded to incorporate directionally-biased transmission in both anterograde (pre to post synptic) or retrograde (post to pre synaptic) directions. This study of directionality is novel in that it does not depend on transmission along a single projection, but incorporates directional transmission across the brain entire network. We reason that if different tauopathic protein species have different directional biases, then perhaps each directional transmission model will differentially fit the spatial patterning of tau distribution from various transgenic mouse models. Specifically, we test a) whether directional transmission fits tauopathy datasets better than non-directional transmission; b) whether we can group tauopathies in mouse models based on our quantitatively defined measure of directional propensity of spread at the macroscopic, whole brain scale, and whether this corresponds with any meaningful disease categories.

We show directional DNT modeling, in the case of some tauopathy datasets, adds more predictive value relative to undirected NT modeling, confirming a role for directionally biased transsynaptic tauopathy expansion. We next demonstrate that retrogradely-biased DNT better recapitulates tau pathology in AD-Like (ADL, here meaning amyloid comorbid; see **Methods:** ***Mouse Tauopathy Datasets*** for further information) mouse models than undirected NT, but a similar strong directional bias is not observed in Non-AD-Like (NADL) mouse models. We tested several other ways of segregating our mouse data using various molecular or etiologic tau-related properties, but none showed reliable differences in spatiotemporal tau spread patterns.

Finally, we explored whether directional network sinks predicted by graph theory can explain why regions such as the EC consistently exhibit earliest pathology in AD and similar conditions. We demonstrate that the network sinks of transregional retrograde tau pathology transmission accurately implicate the EC and other early-tau regions. This suggests regions exhibiting early tau vulnerability might do so due to their location as retrograde network sinks of the retrograde connectivity network, rather than because they act as genuine tau pathology seeds.

## Methods

The data in the present study for creating the mouse connectivity network comes from the Allen Institute for Brain Science’s Mouse Connectivity Atlas (MCA). This network is derived from viral tracing studies and contains fully directional connectivity intensity information from 426 regions across both hemispheres; more information on the MCA can be found at the Allen Institute’s website and in the citation (Oh, et.al., 2014). Regional tau pathology data came from a group of papers all written within the last 10 years, all using IHC for quantification or semi-quantification, and all using mice with a P301S mutant human tau background. All modeling was done using standard Network Transmission (ie, Graph Diffusion; NT), Directed Network Transmission (DNT), and Spatial Diffusion (SPD). For visual examples of the analyses, metrics, and models employed by this study, please refer to Figure 1.

**Figure 1.**
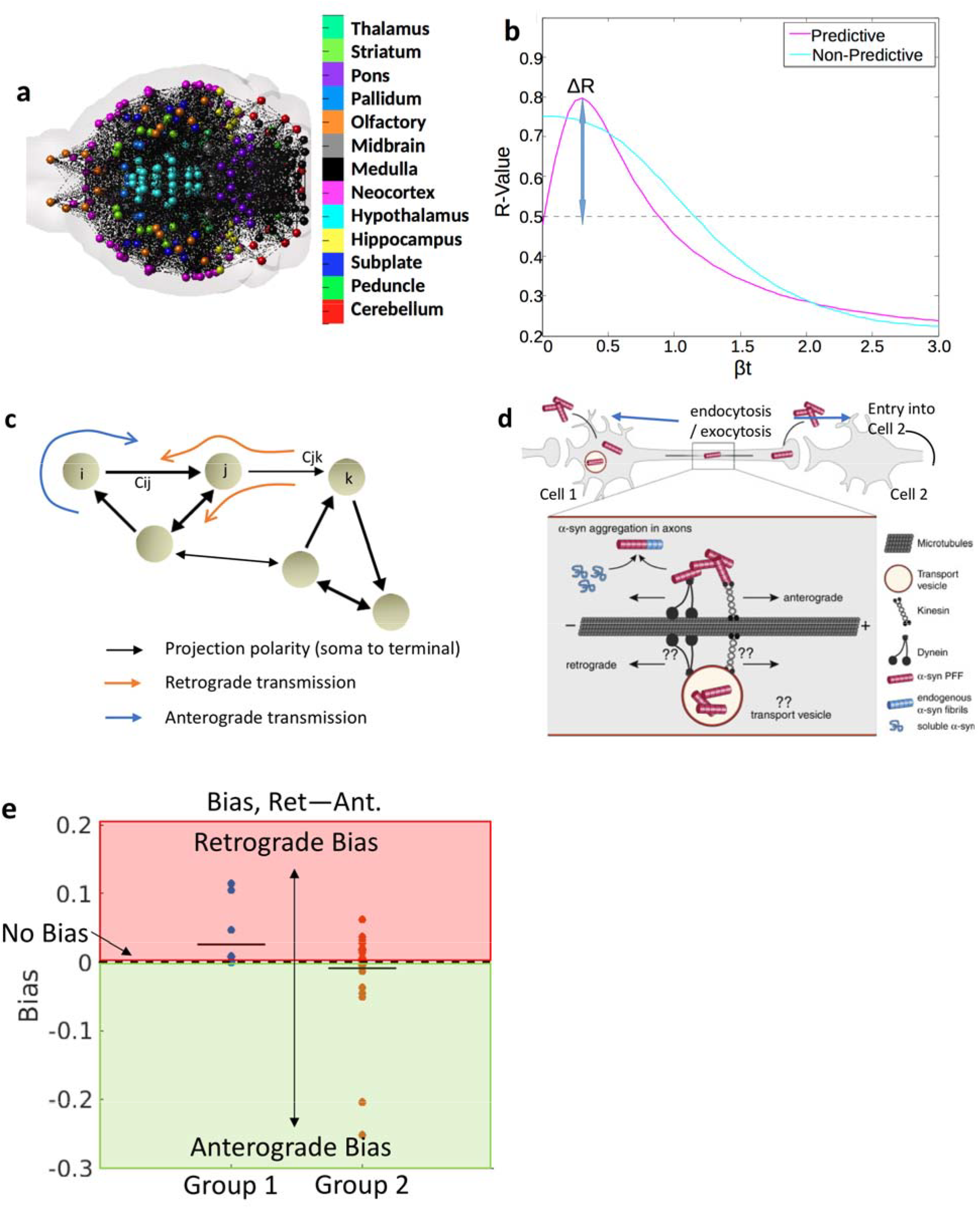
An example figure for the concept of networks, βt-curves, and directional bias. (a) An illustration of the mouse brain network, with balls representing regions and black lines representing connections. (b) An example βt-curve, with the max highlighted, a visual explanation of predictive and non-predictive curves, and the ΔR difference between βt = 0 and βt at max calculation illustrated. (c) Here we show what we mean by directional bias, calculated as ΔR(retrograde) – ΔR(undirected), where red = bias away from retrograde, green = retrograde bias, and the mean of each example group is show as a sold black line.

### Mouse tauopathy datasets

All datasets used in this study had to meet the following criteria: they had to be 10 or fewer years old, they had to use IHC as the basis for all quantifications or semi-quantifications, the tauopathic genetic background of the mice had to be P301S mutant human tau or wild-type human tau to avoid conflating background mutant tau protein type with other analyses, and at least 10 separate brain regions had to be quantified for tau pathology. The datasets publicly available meeting the above standards were as follows: Iba, et.al., 2015 injected P301S mice with P301S tau fibrils and quantified regional pathology in 156 areas at 1 month, 3 months, and 6 months post-injection. Iba, et.al., 2013 injected P301S tau mice with tau fibrils cloned from reverse engineered tau cDNA from misfolded tau isoforms found in AD and injected them into the hippocampus and striatum of P301S mice, generating two separate datasets of 138 regions each. Boluda, et.al., 2015 injected brain homogenate from post mortem confirmed cases of DSAD (with post mortem confirmation of amyloidopathy comorbidity) and CBD into P301S mouse hippocampi, creating two datasets of 106 and 82 quantified regions, respectively. Both Boluda et.al., 2015 and Iba, et.al., 2013 quantified the same timepoints post injection as in Iba, et.al., 2013. Hurtado, et.al., 2010 used mice double transgenic for APP with the Swedish mutation and the P301S tau mutation and spatiotemporally staged their endogenously developing (non-injected) tauopathy at 4 timepoints (2, 4, 6, and 8 months post-birth) across 45 regions. Ahmed, et.al., 2014 injected P301S mice with P301S tau fibrils into the dentate gyrus and visual cortex; from this research we use their published work (20 regions, 4 timepoints). Clavaguera, et al., 2009 injected ALZ-17 mice, which express the human WT version of the longest 4R tau isoform, with P301S tau extracted from aged P301S transgenic mouse model brain homogenate, quantifying 11 regions per hemisphere across 3 timepoints of 6, 9, and 15 months post-seeding. Finally, we use 7 datasets from Kaufman, et al., 2016, which injected different pathological tau strains into the hippocampus of mice and all semi-quantified pathology in 46 regions at 3 timepoints. Further information about each dataset or study used in the present article can be found at each set’s respective citation.

Next we describe how we extracted data from data tables, scatterplots, and quantitative or semi-quantitative anatomical mappings for each dataset. For data derived from Boluda, et al., 2015, Iba, et al., 2013, Iba, et al., 2015, and Kaufman, et al., 2016, data had to be manually extracted from heat map figures displaying semi-quantified pathology anatomically. Using the scaling gradations available in these figures to roughly recreate the semi-quantitative range and steps, and using the Allen Brain Institute’s mouse reference atlas (url: https://mouse.brain-map.org/static/atlas) as an anatomic reference, the authors performed rough remappings into ABI space of the semi-quantification across all quantified brain regions. Data from Clavaguera, et al., 2009 and Ahmed, et al., 2014 were derived from main text data tables or scatterplots, allowing for more precise reconstruction of their quantifications. Data from Hurtado, et al., 2010 represents disease staging per region, more than quantified or semi-quantified regional pathology, but was derived from a supplementary data table.

We additionally split the above cited studies into groupings based on the following criteria: those using ADL tau and those using NADL tau, those using tau derived from human or nonhuman sources, those using P301S tau as the base, and if the tau was exogenously seeded whether the tau was of synthetic or non-synthetic origin. To further characterize ADL and NADL tau, ADL tau simply means that, in the case of exogenous tau inoculation, the injected tau was derived from human AD patients. In the case of endogenous tau pathology models, they all were double or triple transgenic mouse strains having not only a tau mutation but also an amyloid and/or Psen mutation.

### Connectivity metrics, NT, and DNT

Connectivity metrics are calculated based on the topology of the mouse brain network derived from the MCA, where connectivity is represented as “outgoing” along the rows and “incoming” along the columns (Oh, et.al., 2014). Generating a directed and weighted connectivity measure with seed regions was therefore done by summing the weighted row-wise or column-wise values in the MCA at the seed nodes. We could then correlate each selected region’s measured tau pathology with its weighted connectivity, in both anterograde and retrograde directions, with the seed nodes in each study. Further information on using the MCA and its connectivity data for graph analyses can be found at the original citation (Oh, et.al., 2014) and in our prior paper using this atlas (Mezias, et al., 2017).

NT and DNT modeling is similarly dependent on the MCA in this study, and on regional graph adjacency to the seed nodes or nodes already containing pathology. We first calculate the Normalized Adjacency or Laplacian Matrix, *L*, given by the equation below:

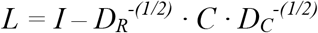

where *C* is the connectivity matrix from the MCA and *D*_*R*_ and *D*_*c*_ are the row and column wise diagonal matrices from the MCA. Previous graph theoretic work has characterized diffusion over a network from defined seedpoints and has been shown to be predictive for both volumetric loss (Raj, et.al., 2012) and metabolic dysfunction (Raj, et.al., 2015) in patients with AD or other tauopathic dementia, as well as for mouse models of tauopathy and amyloidopathy (Mezias, LoCastro, & Raj, 2016):

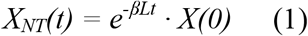

Here *L* is the Laplacian Matrix, β*t* is a constant representing diffusion time determined by when it maximizes ΔR, which is the change in r-value from β*t* = 0 to maximum, *X(0)* is a vector with *n* number of nodes length representing seeding, and *X(t)* is a vector with *n* number of nodes length representing the final pattern of network diffusion. For a visual example of β*t*-value parameter optimization in this model, pease refer to Figure 1b. Further information on the derivation of the above equation and graph diffusion modeling can be found at the aforementioned citations.

In the present study we place the NT model equation inside an iterative process so that we both capture the directionality information from the MCA and so that represent tauopathy as not only a process of spread but also of accumulation, analogous to a prion-like disorder. The iterative algorithm over timesteps, in our case months, works in the following manner:

1. Initiate the NT model using *X(0)*, the seed or baseline pathology
2. Calculate new regional pathology values for that timestep
3. Re-initialize the NT model for the next timestep using the new regional pathology values in (2) above as the ‘seed’ or ‘baseline.’ Repeat this process until *t(i)* == *t(n)*.

We capture the directionality information from the MCA by calculating different networks for each direction, where *C* represents the standard connectivity network, *C*^*T*^ represents the network in the retrograde direction, and the summed connectivity of both, *C* + *C*^*T*^, represents the undirected network derived from the MCA. Our overall approach using the iterative algorithm described above, as well as capturing directional networks, can be summarized with the following equation:

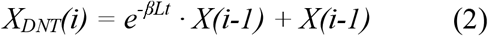

The summative approach implemented approach in 2 arrives at similar results, in terms of R-value, to the standard Network Diffusion model, but better represents spreading and accumulating tau spread physiologically. We can therefore interpret our implementation of an accumulative graph diffusion model as the integral of the standard network transmission model in Eqn. 1 above, with seeds changing per iteration, or in the case of the integral, per round of spread of the diffusive tau. More information on the mathematics behind this approach can be found in our previous work on the subject (Mezias, et al., 2017).

### DNT Bias Calculations and a Proportional DNT Model

To test for potential directional spread biases, we created a metric to assess bias between modeled purely anterograde and purely retrograde spread as well as a DNT model allowing for proportionate spread in each direction, determined by a single parameter. For visual examples of directional transmission and what is meant by anterograde or retrograde spread, please refer to Figures 1c & d. For a visual example of how we present directional bias, please refer to Figure 1e.

1. **Directional Bias.** To determine directional bias, we performed a simple calculation of subtracting ΔR-values from anterograde DNT from retrograde DNT ΔR-values. A value of 0 would indicate no bias, a value >1 would indicate a bias in the tested direction, and a value <1 would indicate a bias away from the tested direction. The above cited calculations all follow the form of this example equation:

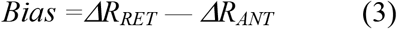 We additionally calculated bias relative to modeled non-directed spread by subtracting either anterograde or retrograde ΔR-values from undirected NT ΔR-values. This calculation follows the same form as Eqn. (3) above.
2. **Proportionate DNT Model.** Our proportionate DNT model is designed to model spread occurring simultaneously in both the anterograde and retrograde directions, but in different proportions. We choose the proportion via a parameter we will here call the s-value, which we modulate between 0 and 1. The proportionate DNT model is the same as the standard DNT model from Eqn. (2) in the prior section with one important step added to the process as we first modify the connectome via the following equation:

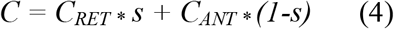 S-values must always be between 0 and 1. In all cases, in this current paper, we modulate s-values between 0 and 1 by steps of 0.05. We define peak s-value as the one which produces the peak r-value.

### Data analysis and statistics

All analyses performed were a variant of a standard linear regression using natural log transformed regressions due to empirical pathology and predicted values being best fit by exponential distributions. We first compared regional gene expression with DNT to see which method added more predictive value in addition to undirected NT, using methodology similar to a prior paper from our group (Mezias, et al., 2017). We next examined whether tau pathology preferred to spread in anterograde or retrograde directions using this bias and proportionate DNT calculations from Eqns. (3) and (4), respectively. We compared across groupings of tau pathology (ADL vs. NADL; human vs. nonhuman origin; P301S vs. non-P301S; synthetic fibril tau origin vs. non-synthetic origin) both by comparing, for example, ADL and NADL bias and peak s-value distributions using two-sample t-tests, and by comparing, for instance, ADL tau bias or peak s-value distributions to a simulated normal distribution centered around 0 using a one-sample t-test. To test for the effects of progression time on directional bias in tau propagation we correlated each measured timepoint’s retrograde and anterograde bias, across all studies, and correlated that with the time of measurement (in months) after study initiation. All statistical analyses were performed using MatLab.

### Random and whole brain seeding simulations

Several graph theoretic simulations were performed to further understand the network basis of observed selective vulnerability of certain brain regions as the earliest seeds of tau pathology.

1. **Random seeding**. First we build null random models of DNT after repeatedly seeding randomly a single or small subset of regions, in order to assess whether specific seeding in a particular region or subset of GM areas was a necessary precondition for observing AD like pathology in a graph diffusion model. At each run of the DNT, a random seeding was initiated, such that 0.5% to 10% of GM nodes in the mouse brain connectivity network were randomly chosen to be “seeds” for simulated tauopathy. The random seeding simulations were run in two different manners: a) 1000 random seeding simulations were run through 2 or 8 iterations and the regions of highest pathology using anterograde or retrograde DNT, or undirected NT were recorded; and b) from these 1000 random seeding simulations 25 were randomly selected and their regional pathology levels were averaged using a standard bootstrapping procedure to generate averages (Efron, 1979). We then correlated the DNT prediction following these random seeding simulations using anterograde, retrograde or undirected NT, with data from our non-exogenously seeded mouse AD-Like mouse dataset, Hurtado, et al., 2010. We ran all random seeding simulations twice to generate the 3D scatterplots and brain illustrations, respectively, in Figure 6.
2. **Whole brain diffuse seeding simulations**. This refers to the application of DNT with initiation with equal weight throughout the whole brain (all 426 ABA regions) simultaneously. The rest of the analysis was as in #1 above.
3. **1**^**st**^ **Eigenvector evaluation**. It was previously conjectured that the eigenmodes of the brain’s connectivity network could predict spatiotemporal degenerative pathology progression (Raj, et al., 2012). We therefore subjected the mouse brain connectomes (anterograde, retrograde, undirected) to an eigenvalue decomposition and used the first eigenvector as a predictor of the expected regional tau pathology. This eigenvector predictor was statistically compared to empirical regional tau pathology, as above.

## Results

### Directionality in connectivity networks improves connectomics based modeling of tau pathology

Recent prior research indicates that, in exogenously seeded mouse models, connectivity with afflicted regions drives subsequent regional tau pathology severity and spatiotemporal tau progression more so than gene expression similarity with regions already affected. Expression levels of tauopathic or degenerative disease ‘risk-factor’ genes also do not predict observed tau pathology patterns or their spread over the brain’s connectivity network (Mezias, et al., 2017). However, that prior research, along with other previous work modeling tauopathy spread, is limited by using only undirectred connectomes. Whether tau pathology spread can be directionally biased relative to axonal polarity (ie, anterogradely or retrogradely biased) is a presently unanswered question. Here we explicitly test whether employing directional mouse brain connectivity networks improves upon using more typical undirected networks, with tau spread modeled using DNT across the 6 datasets used in Mezias, et al., 2017.

We find that DNT improves on the predictions of undirected NT in 4 out of 6 cases, when compared against the empirical pattern of tau pathology progression (Table 1). This establishes the proposition that directional (whether anterograde or retrograde) transmission of tau can perform better at capturing the observed tau pattern in mouse models compared to undirected transmission.

**Table 1.** Adding directionality to NT models can improve upon undirected NT predictions, but regional gene expression information does not. In the above table we compare ΔR-value predictions from our undirected NT model, as compared with anterograde and retrograde DNT, as well as with transmission based upon regions with similar genetic profiles. All modeling of pathology spread has proteinopathy initiating at the reported seedpoints, enumerated in the table above, from each study. The largest ΔR-value, per study (per row), comparing across diffusion models, is bolded.

| Study | Injection Region | Injectate | # Regions | NT-NoDir. | DNT-Ant. | DNT-Ret. |
| --- | --- | --- | --- | --- | --- | --- |
| Boluda, et al., 2015 | Hipp. | DSAD homogenate | 102 | 0.29 | 0.05 | **0.30** |
| Boluda, et al., 2015 | Hipp. | CBD homogenate | 94 | **0.23** | 0.00 | 0.22 |
| Iba, et al., 2013 | Hipp. | Synthetic Fibrils | 96 | **0.45** | 0.32 | 0.37 |
| Iba, et al., 2013 | Striatum | Synthetic Fibrils | 76 | 0.21 | 0.12 | **0.39** |
| Clavaguera, et al., 2009 | Hipp. | P301S tau | 22 | 0.22 | 0.18 | **0.30** |
| Hurtado, et al., 2010 | N/A | N/A | 45 | 0.02 | 0.00 | **0.13** |

### ADL mouse models show a bias towards retrograde tau transmission

We next tested whether directional preference is specific to certain molecular or etiologic properties in studies mouse models. We first assess whether tau being ADL, defined as tauopathy that became pathological comorbid with amyloidopathy, influenced spread patterns (see **Methods:** ***Mouse Tauopathy Datasets*** for further information). Figure 2a shows that regional tau pathology severity and spatiotemporal progression patterns of 75% of ADL studies (as defined in Introduction and methods) are best recapitulated by retrograde DNT. Anterograde DNT fits all ADL datasets poorly. This is true whether we seeded DNT at the reported exogenous seeding location or using the baseline pathology pattern. In contrast, the picture is much more mixed as to which model best recreates NADL tau pathology patterns (Figure 2a), with some datasets favoring anterograde, some retrograde and others undirected transmission.

**Figure 2.**
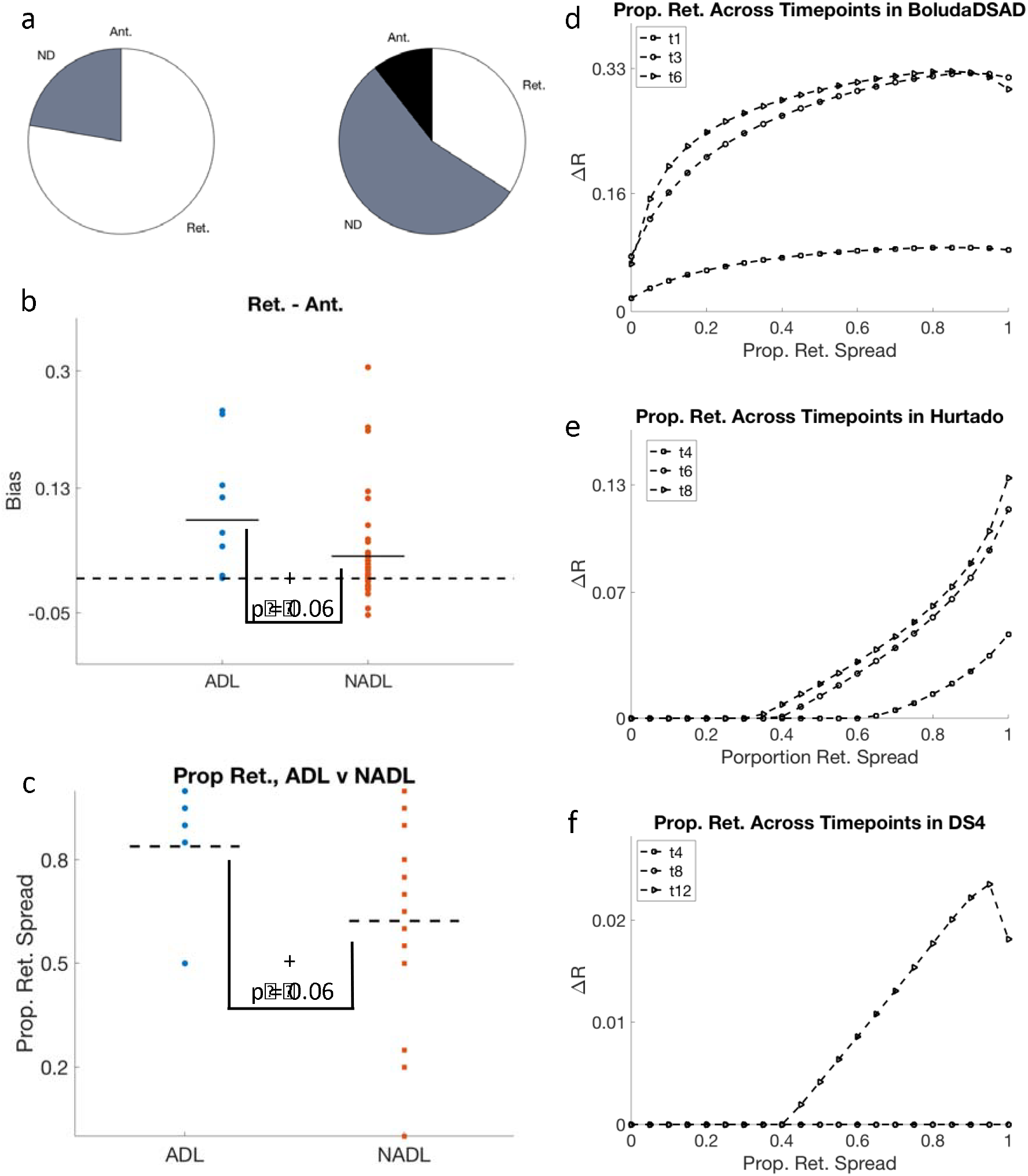
ADL tau shows a consistent retrograde bias in its spread while NADL tau does not. (a) Retrograde DNT was the best model in 75% of measured timepoints across all ADL tau studies. NADL tau studies showed no such directional biases, with the DNT or undirected NT model best recreating tau pathology in these studies being split between all models fairly evenly. (b) AD-Like (Amyloid Comorbid) tau pathology, across studies and measured timepoints, demonstrates a retrograde spread bias relative to both Non-AD-Like tau and null hypothesis of no bias, when DNT modeling starts from reported seedpoint. (c) ADL studies have higher proportions of retrograde spread producing the best ΔR and R-values relative to both the NADL group of studies, and relative to a null hypothesis of a 0.5 proportion of retrograde spread (fully bidirectional). All ADL datasets, obtained from Boluda, et al., 2015 (d), Hurtado, et al., 2010 (e), and Kauffman, et al., 2016 (f) show a strong bias towards better fitting at higher proportions pof retrograde spread. The asterisks and crosses above a distribution indicate that the distribution is significantly different from 0 in a one-sample t-test and the asterisks and crosses below the brackets hanging downwards between each pair of distributions indicates the distributions are significantly different from each other in a two-sample t-test. * p < 0.05, + p < 0.10.

We next calculate the metric for measuring directional bias, *ΔR*_dir_, between anterograde and retrograde spread by subtracting the ΔR using anterograde DNT from the ΔR using retrograde DNT (See Eqn. 3 in **Methods**). In our metric, a positive bias number indicates retrograde driven tau pathology spread, while a negative number indicates anterograde driven progression. Initiating DNT modeling from baseline pathology measurements across studies shows ADL tau studies trend towards being significantly more retrograde in their pathology development than NADL studies using a two-sample t-test, t = 1.90, p = 0.06, and trend towards being significantly more retrograde than a null hypothesis of no directional bias using a one-sample t-test, t = 1.86, p = 0.06 (Figure 2b). We obtained no other significant results for any other tested differentiating groupings of tau pathology types used across studies (S.Fig. 3a-b).

**Figure 3.**
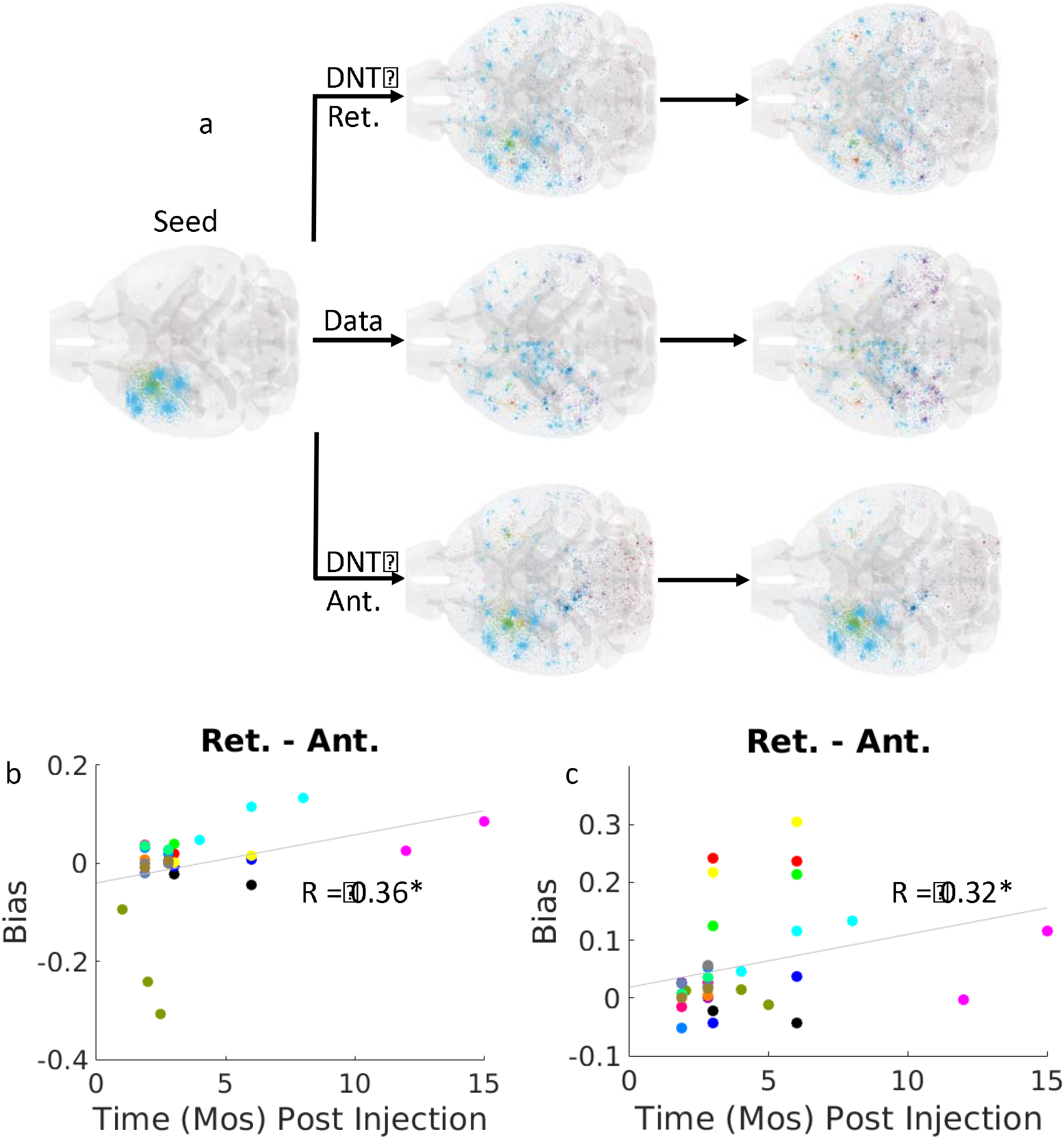
Tau pathology exhibits, across studies, a progressive shift towards retrograde spread. In (a) we illustrate Iba, et al., 2013, modeled by retrograde and anterograde DNT, as compared with the empirical spatiotemporal tau pathology progression, as an anatomical demonstration of panels b & c below. From both baseline pathology measurements (b) and reported seedpoint (c), collapsing across studies and timepoints, we found pathology measured at later timepoints became progressively better recapitulated using retrograde DNT as compared with anterograde DNT (or undirected NT). In b & c each color represents a study, so studies can be tracked over time in these panels. * p < 0.05, ** p < 0.01.

We also tested a bias measure relative to non-directed NT. ADL studies exhibit either a significant or trending toward significant bias towards retrograde spread as compared with undirected spread using two-sample t-tests, and as compared with the null hypothesis of no directional bias using a one-sample t-test, whether modeling is begun at reported seedpoint (S.Fig. 3c) or baseline pathology measurement (S.Fig. 3d). Mice injected with human origin tau also show a significant bias away from anterograde spread, but no other grouping (P301S versus non-P301S mouse background; synthetic or non-synthetic origin of tau) produced results that were significant across these analyses. Individual study fitting curves for regional pathology rate of increase patterns can be found in Figure S1 for DNT beginning at reported seedpoint, and in Figure S2 for DNT initiating using baseline tau pathology.

### Retrograde biased porportional DNT models are uniformly better predictors of ADL, but not consistently NADL, tau pathology datasets

Employing the proportionate DNT model described in **Methods** (See Eqn. 4), we tested whether a higher proportion of spread being retrograde would lead to a better fit with ADL datasets, here measured with R and p-values, and whether any bias could be observed in NADL datasets. Across all 3 ADL datasets, we find that proportionate DNT models work best when the retrograde direction accounts for the large majority of tau spread. When we take only the proportion of retrograde spread best recapitulating each timepoint of each study, we find that our best modeled propagation is 0.85, or 85% retrograde, for ADL datasets, but is 0.625, or 62.5% retrograde, for NADL datasets (Figure 2c). In a two sample T-test, we find the proportions of spread of best fit for ADL datasets trend towards significantly more retrograde than those of NADL datasets, T = 1.91, p = 0.06 (Figure 2c). In a one-sample T-test we find that the distribution of proportions of retrograde spread among ADL datasets is significantly more retrograde than a null-hypothesis of spread being equally bidirectional where proportion of retrograde spread is 0.5; T = 2.01, p = 0.04 (Figure 2c).

We finally verify that these results are not due to a single outlier dataset or timepoint. In an ADL dataset from Boluda, et al., 2015 using tau from DSAD patients, the proportion of retrograde spread for the peak perfoming DNT model never falls below 0.85 at any point (Figure 2d). In another ADL set using a non-seeded APPswe x P301S genetic mouse model from Hurtado, et al., 2010, the proportion never falls below 1.00 (Figure 2e). Unexpectedly, in a dataset obtained from Kauffman, et al., 2016 using purified tau strains, at very early timepoints we find no good fit with the data at early timepoints regardless of whether spread is biased proportionally towards reotrgrade or antergroade spread. However, these timepoints are both prior to 2-months following tau injection into the mouse; by the final timepoint of 3 months post-injection, a proportion of retrograde spread of 0.95 or 95% retrograde best recreates the data (Figure 2f). We see no consistent pattern of any directional bias across NADL datasets. Individual NADL dataset visualizations such as those for ADL sets in Figure 3d-f can be found in Figure S.4.

### As tau pathology progresses its spread becomes increasingly driven by retrograde connectivity based transmission

We next test whether the duration of tau pathology affects its spread pattern. Across all studies, we found that at later time points tau pathology becomes increasingly well modeled by retrograde DNT as compared with anterograde DNT, regardless of whether pathology modeling was initiated at reported seedpoint, r = 0.36, p < 0.05 (Figure 3b), or baseline tau pathology measurement, r = 0.32, p < 0.05 (Figure 3c). We illustrate this anatomically in Figure 3a, using the striatal injection data from Iba, et al., 2013 as an example. Our results accordingly indicate an increasing bias towards retrograde tau pathology spread and away from anterograde tau pathology spread as the elapsed time since the initiation of tau pathology increases. We found a similar, albeit non-significant pattern when we just considered direct connectivity with the seed region: ADL tau pathology was, over time, better predicted by retrograde connectivity than by anterograde connectivity, but there was no such bias for NADL tau (Fig. S5).

### Progressively retrograde tau pathology spread biases are stronger in ADL datasets, but persist even in NADL datasets

The correlation between timepoint of measurement and degree of bias towards retrograde spread in ADL studies is significant, p < 0.01, and is stronger than the same above reported effect across all studies, r = 0.67 (Figure 4a). However, the same pattern of progressive retrograde spread biases in tau pathology over time emerges in NADL studies, r = 0.28, albeit it is non-significant, p = 0.13 (Figure 4b). We further tested which proportion of retrograde spread produced peak r-value between our proportionate DNT model and the datasets, across timepoints. The proportion retrograde spread was significantly positively correlated with timepoint of measurement in ADL datasets, r = 0.63, p < 0.01, indicating a higher proportion of tau pathology spread being retrograde over time (Figure 4c). Akin to our results using directional bias above, NADL datasets exhibited a much weaker and nonsignificant, albeit positive relationship between proportion retrograde spread and time, r = 0.18, p = 0.21 (Figure 4d). This effect cannot simply be due to tau spread speed differences between ADL and NADL dataset groups, as our model constant for diffusion speed, β, is not significantly different on average between ADL and NADL groups (Figure 4e). We further find that among ADL studies the distribution of best-fit 100% anterograde or 100% retrograde DNT β values are not significantly different either (Figure 4f). We therefore demonstrate tau spread becomes even more biased towards retrograde propagation in ADL studies than NADL studies for reasons unrelated to the speed of pathology diffusion. However, we remained unconvinced that the exhibited progressive retrograde bias in tau pathology propagation was due to whether a dataset was ADL or NADL, given the results from Figure 3, reported in the prior section.

**Figure 4.**
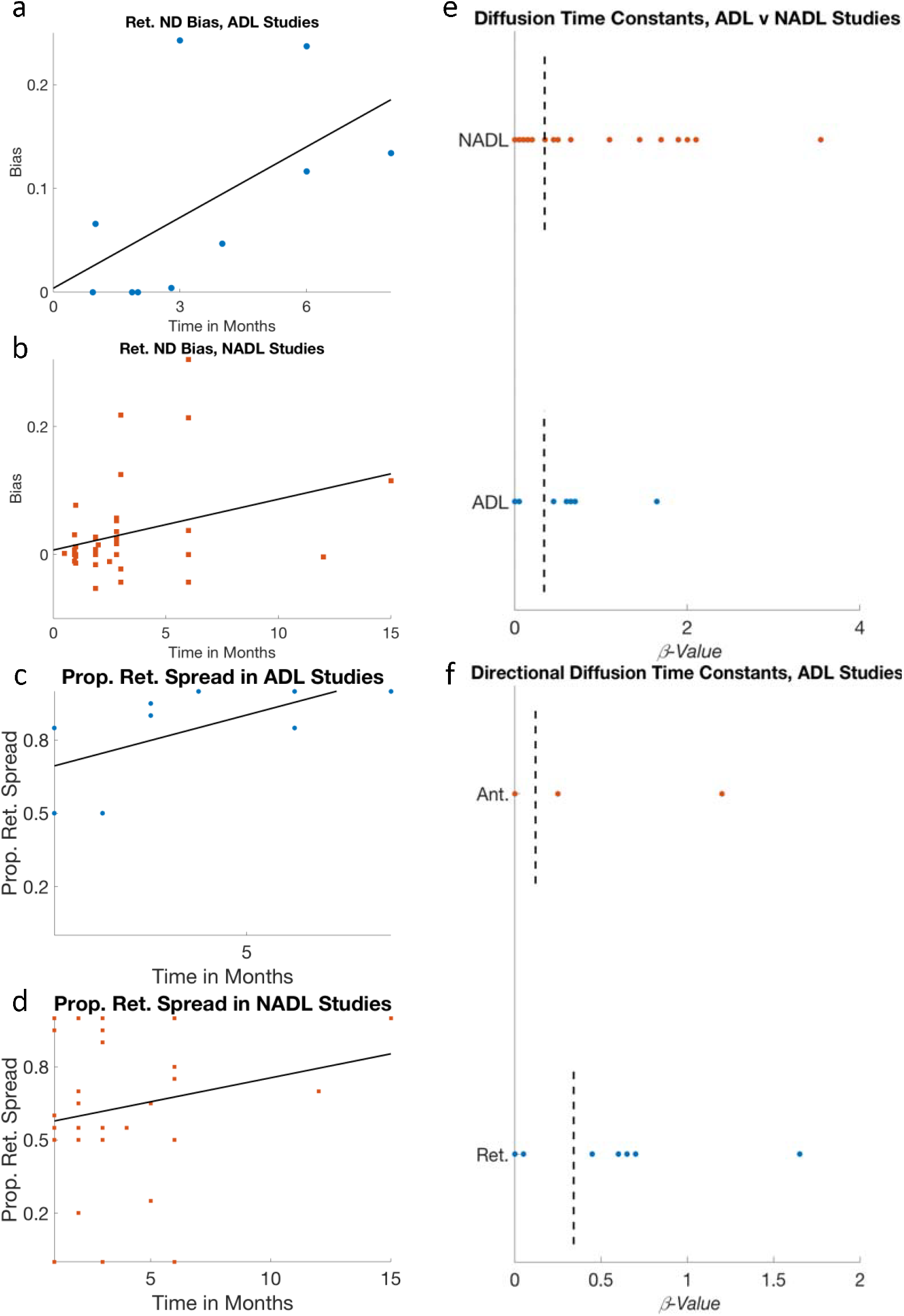
The progressive development of a retrograde spread bias in tau pathology cannot be explained alone by each dataset’s ADL status. (a) ADL datasets exhibit an especially strong development of a retrograde bias in tau pathology spread. (b) While this same progressive development of a retrograde bias in NADL studies is non-significant, the same pattern exists as in ADL tau datasets. Similarly, when we run a spectrum of proportion of retrograde spread from fully anterograde, prop = 0, to fully retrograde, prop = 1, we find that (c) ADL studies show a significantly increasing biases over time towards higher proportions of retrograde spread, while (d) NADL studies show a similar but weaker and nonsignificant pattern of results. (e) This difference between NADL and ADL datasets in their degree of development of a progressive retrograde bias in tau pathology spread cannot be explained by differences in diffusion speed alone as the β diffusion speed constant values of best fit are the same between the two groups. (f) Distributions of β diffusion speed constants are the same, across all ADL studies, whether 100% retrograde or 100% anterograde DNT are used, underscoring the results in panel (e). * p < 0.05, ** p < 0.01.

### Vulnerable sites in AD: pathology initiators or pathology accumulators?

The above evidence for retrograde bias in tau transmission opens the possibility that the early sites of tau pathology (EC, retrosplenial, temporal cortex) might be better described as retrograde accumulators of diffuse pathology rather than initiators of pathology. To begin with, network “sinks” can be mathematically defined by the connectivity matrix’s “eigenmodes”, especially the first one, which represents the long-term steady-state pattern of spread on the network, regardless of its starting point (Raj et al., 2012). The network given by matrix C can be decomposed into its constituent “eigenvectors” *U* = {*u*_*i*_,*i* = 1: *N*_*rois*_}, where *LCUΛU*^*T*^ is the eigen-decomposition. The first eigenvector, *u*_1_, can point to nodes or regions that are a network “attractor” into which any diffusing entity will preferentially accumulate. We therefore hypothesized that *u*_1_ of the retrograde network Laplacian would produce the highest values in entries associated with regions of early tau pathology. In Figure 5a we show a 3D scatterplot and anatomical illustrations of regional patterns predicted by the first Laplacian eigenvector *u*_1_ evaluated in retrograde and anterograde networks. The pattern given by *u*_1_ (Figure 5a) correctly predicts which regions are the most important accumulators of AD tau pathology, with the EC, piriform and retrosplenial areas having the highest values. In contrast, *u*_1_ of the anterograde network Laplacian has high values in striatal and midbrain structures, which are not involved in early AD tauopathy (Figure 5a).

**Figure 5.**
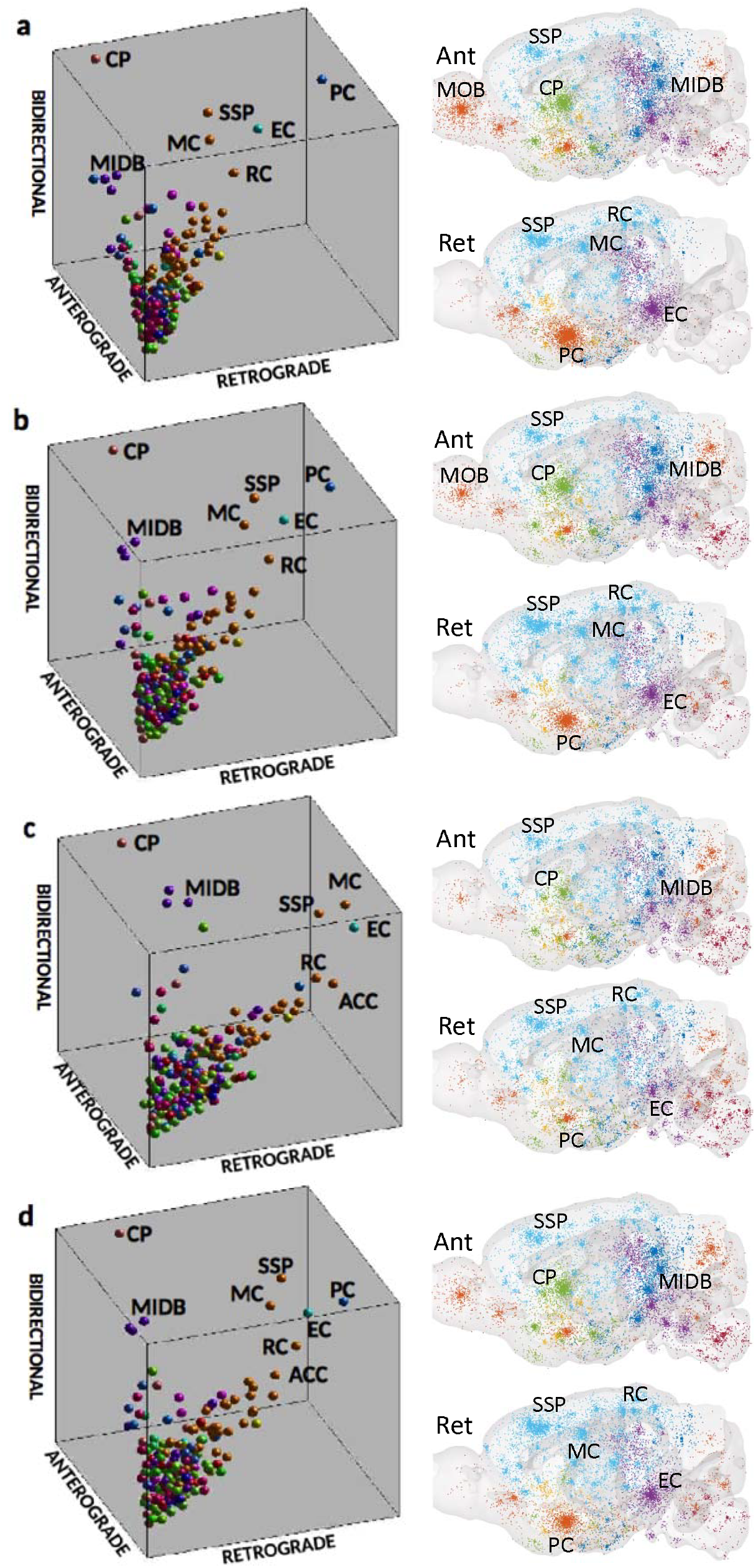
Theoretical random seeding, whole brain simultaneous pathology initiation, and *u1* using retrograde DNT select regions known to be exhibit early and outsized tangle pathology as those most liekly to be tau proteinopathy accumulators. In the 3D scatterplots and mouse brain illustrations above, when transmission proceeds in the retrograde direction, (a) *u1,* (b) whole brain simultaneous seeding, (c) and random pathology initiation with long and (d) short diffusion times all implicate regions such as the EC, ACC, RC, and PC, areas known to exhibit early and heavy tau pathology, as the regions most liklely to accumulate tauopathy here.

### Focal seeding is not necessary to see pathology accumulation in regions associated with early tauopathy in AD

Using random and diffuse seeding simulations, we study whether tau accumulation into areas of early tau pathology can come about from random or diffuse pathology seeding. We hypothesized DNT-driven accumulation from diffuse or random sources would implicate regions commonly associated with early AD tangle pathology, e.g. the EC. In Figure 5 we show 3D scatterplots and anatomical illustrations of regional patterns predicted by various ways of applying the DNT model following: (1) whole brain simultaneous seeding, (2) random pathology initiation with long (8 iteration) and (3) short (2 iterations) diffusion times. Limbic areas are prominent in the retrograde DNT simulations, whereas striatal areas are prominent in the anterograde DNT simulations. Indeed, the most prominent accumulators of pathology, as predicted by retrograde DNT after both diffuse whole brain (Figure 5b) and random (Figure 5c & d) seeding, include: EC, retrosplenial, and piriform cortices, all regions associated with early tangle burden in AD (Greicius, et.al., 2004). Neocortical motor and somatosensory regions were also implicated (Figure 5a-d). When DNT was run through only two iterations of DNT after random seeding, representing a very early stage of tangle pathology, the EC was the most prominent accumulator of tauopathy (Figure 5c).

Finally, we test whether random seeding, on average, can lead to accumulation into the sites of early tau pathology. Retrograde DNT was run on the mouse connectome over 1000 random seeding simulations. We found that random seeding with subsequent retrograde DNT produced a large range of strong correlations with empirical spatiotemporal tau pathology pattern observed in our non-exogenously seeded ADL mouse tauopathy dataset (Figure 6a-e; Hurtado, et al., 2010). This was true across all timepoints, and the resulting Pearosn’s r-statistic was comparable to those generated using retrograde DNT following seeding with the initial reported pathology pattern, especially at later timepoints, median r = −0.03, 0.08, 0.17, 0.22 (S.Fig. 6), median Δr = 0.01, 0.09, 0.18, 0.24 (Figure 6d), at 2, 4, 6, and 8 months, respectively. The median Δr values from random seeding simulations were in line with, or even better than, the Δr values from retrograde DNT initiated at the baseline pathology measurements from Hurtado, et al., 2010 (Figure 6d). Anatomic illustrations of a randomly selected random seeding simulation are found in Figure 6b, with the empirical spatiotemporal tau pathology illustrated in Figure 6a. We also modeled whole brain simultaneous pathology initiation and obtained results similar to but even more striking than those from random tauopathy seeding above, as not only Δr values, but in fact r-values (Figure 6e) between data and modeled pathology were in line with or better than those obtained using the baseline pathology measurement from Hurtado, et al., 2010. Anatomical illustrations of whole brain seeding are in Figure 6c. However, both random seeding and whole brain simultaneous yielded only moderate recreations of empirical spatiotemporal pathology patterns from Hurtado, et al., 2010 (Figure 6; S.Fig. 6). These theoretical results provide further evidence that retrograde transmission is the most likely mode of AD-related tau pathology spread and that early tauopathy sites are more likely accumulators rather than initiators of pathology, purely as a consequence of network topology.

**Figure 6.**
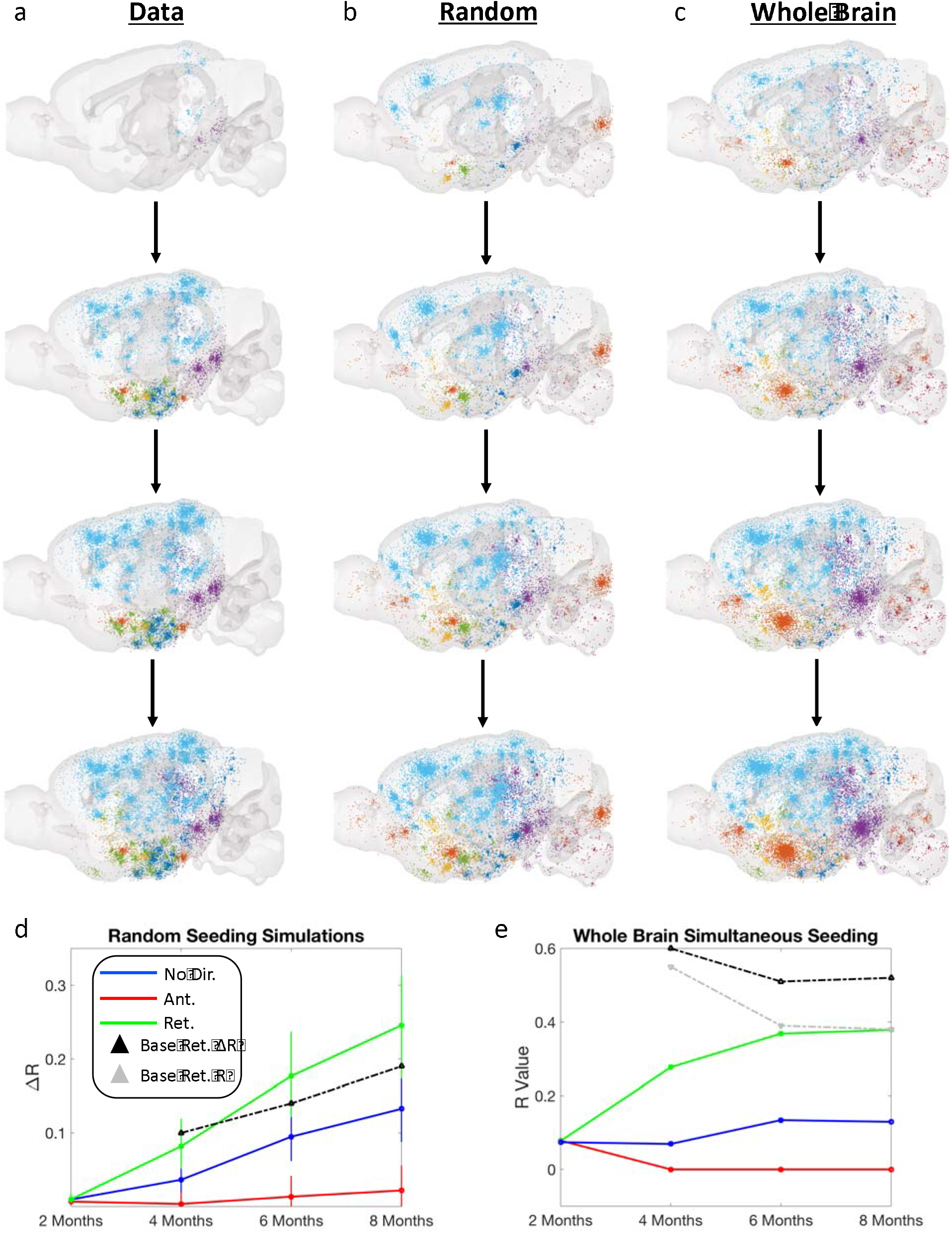
Both random and whole-brain simultaneous seeding simulations followed by retrograde DNT perform well at recreating regional severity and spatiotemporal development of tau pathology in an ADL mouse model without exogenous seeding (Hurtado, et al., 2010). We first anatomically illustrate spatiotemporal tau pathology progression using (a) the empirical data, (b) a single random seeding simulation using retrograde DNT, (c) and the whole brain simultaneous seeding simulation using retrograde DNT. (d) We display, over time, the median Δr-values from all 1000 simulations, in a line plot, per NT/DNT model used, as well as the Δr-value initiated at baseline pathology using retrograde DNT as a comparison. The error bars are plotted at the 25^th^ and 75^th^ percentiles of all 1000 Δr-values, per ND/DNT model used. (e) Here we plot the r-values from whole-brain simultaneous seeding with the data from Hurtado, et al., 2010, per NT/DNT model used, over time, as well as the r-values from baseline pathology using retrograde DNT, and the baseline pathology, as a comparison. Please refer to the legend for interpreting the line plot for more information.

## Discussion

While prior studies firmly established the predominance of transsynaptic spread mechanisms in driving spatiotemporal pathology development (Ahmed, et al., 2014; Clavaguera, et al., 2009; Tai, et al., 2014; Holmes, et al., 2014; Iba, et al., 2015; Iba, et al., 2013; Boluda, et al., 2015; Kaufman, et al., 2016; Raj, et al., 2012; Mezias, et al., 2017), the present research is the first to explore whether the direction of transsynaptic tau spread might be different across conditions. We find ADL tau was best fit by DNT modeling in the retrograde direction, whereas NADL tau pathology (here, derived from FTD or CBD cases and mutations) did not show a retrograde bias. Therefore, ADL and NADL tau must transit transsynaptically via different mechanisms in the brain, even when both start from the same injection site. Furthermore, the current work finds a progressively developing retrograde bias in tau pathology spread, across most of the 17 datasets analyzed here, regardless of whether they are seeded by ADL or NADL tau. Finally, we demonstrate focal seeding in putative pathology initiation regions (EC, LC, hippocampus) is unnecessary to accurately recreate spatiotemporal tau pathology development, working against the SVb hypothesis of spatiotemporal pathology progression. Instead, our simulations suggest that both diffuse and entirely random seeding, followed by retrograde network transmission, yield progressive accumulation in regions usually conceptualized as initiators of AD pathology, prominently, EC, retrosplenial and piriform cortices, more in line with MNx hypotheses of spatiotemporal degenerative proteinopathy progression. Our analyses and their implications are discussed further below.

### Alzheimer’s-Like tau pathology in mouse models exhibits retrograde bias in its spread

Data in Figure 2 indicates ADL tau exhibits a consistent retrograde transmission bias in its spread pattern across all timepoints. 75% of ADL timepoints from all studies were best fit using retrograde DNT, while none were best recreated using anterograde DNT (Figure 2a-b). Furthermore, when we used a proportionate DNT model, ADL models were uniformly best modeled by heavily retrograde spread, whereas on average NADL models appear best modeled by approximately bidirectional spread (Figure 2c-f). This is an important result because it is the first instance, to our knowledge, that a directional difference in the spread of tau pathology between groups of tauopathic disorders has been demonstrated, thus expanding on the hypothetical MNx framework (Warren, et al., 2013) for explaining spatiotemporal pathology differences between tauopathic conditions. What differences might exist between ADL and NADL tau that might explain such a difference in their preferred directions of transsynaptic and transregional spread?

First, tau tangle structures within disease condition are often consistent, while they can vary considerably between disease condition (Clavaguera, et al., 2013). For example, when tau pathology from a human AD patient is injected into a mouse, the structure of tangle aggregates in the mice mirrors that of the patient (Clavaguera, et al., 2013). Pathological tau species have been posited to be able to bypass or move through the axon-soma membrane on an isoform or strain specific basis (Zempel, H., et al., 2017). Thus, it could be that the misfolded tau strains present in AD or ADL tau groups are more readily able to break through or bypass the axon-soma barrier than is NADL tau.

Second, retrograde transmission of tau might be mediated by Aβ, which is known to induce early missorting of tau from axons to dendrites (Ballatore et al., 2007; Fein et al., 2008; Ittner et al., 2010; Takahashi et al., 2010). Within our own datasets, we find circumstantial support for the presence of amyloid indicating a retrograde tau bias: DNT modeling implicated a retrograde bias in tau propagation at even the earliest measurements of proteinopathy in the dataset from Hurtado, et.al., 2010, which was generated using a triple transgenic mouse model comorbid for tau and amyloid. In other studies, intraceerebral Aβ injections in P301L transgenic mice exacerbated hyperphosphorylation of tau and NFT formation, not at somatosensory cortical and hippocampcal injection sites, but rather in the retrogradely-connected basolateral amygdala (Götz, et.al., 2001). This suggests that Aβ induced damage to terminals or axons projecting to the injection site caused NFT formation in presynaptic cell bodies. Such an occurrence would manifest as a retrograde bias in tau pathology spread in the present research.

Some combination of the above explanations is also a viable hypothesis: Aβ induced retrograde tau transmission should appear earlier in the disease course than retrograde spread caused by the breakdown of the axon-soma barrier, as amyloidopathy has been repeatedly shown to precede or co-occur with tauopathy in AD (Bolmont, et.al., 2007; Götz, et.al., 2004; Ittner, et.al., 2010). Regardless, the implications of our findings are that different tauopathic degenerative conditions exhibit divergent spatiotemporal development patterns potentially because their transsynaptic and therefore transregional spread mechanisms differ, expanding upon the proposed MNx framework of degenerative tauopathies.

### Tauopathy propagation might develop a progressive retrograde bias

Retrograde, but not anterograde DNT, was highly predictive of the rate of increase of regional tau tangle burden in longitudinal data (Figure 3). A progressively developing retrograde bias, if confirmed by bench studies, would be a highly novel finding with the potential to alter how we think about tau pathology progression. Therefore, it is instructive to review existing histopathological evidence of retrograde tau transmission emerging from mouse models. Most in vivo and in vitro studies indicate at least a bidirectional transmission of tau (Wu, et.al., 2013; Dujardin, et.al., 2014; Tai, et.al., 2014; Iba, et.al., 2013). Iba et.al., 2013 demonstrated the classic spread of preformed tau fibrils in P301S mouse model, and found that hippocampal seeding yields spread into EC and nearby cortices, raising the possibility that hippocampal efferents were involved. Retrograde DNT best fits end-stage regional tau deposition and rates of increase across studies (Figure 3) and these results suggest that retrograde transmission biases develop progressively.

We advance two potential explanations that build upon the hypothetical MNx framework of degenerative diseases (Warren, et al., 2013). First, an increasing retrograde bias in tau pathology spread may be caused by progressive breakdown of the axon-soma barrier which in the normal state limits retrograde misplacement of tau into somatodendritic compartment. Over the course of tauopathic disease, repeated exposure to hyperphosphorylated tau erodes the barrier, leading to increasing levels of retrograde missorting of tau from axon to soma and dendrites (Li, et.al., 2011). However, it could also be that the composition of strains of misfolded tau, over time, become increasingly dominated by those biased towards retrograde mecahnisms of transsynaptic transport. There is some basis in literature for this hypothesis: first, that structurally distinct misfolded and pathological strains of tau exist has been confirmed by bench research (Kaufman, et al., 2016). Furthermore, structural differences between gross NFT structure between disease conditions, such as between AD and FTD, that remain consistent with disorder type, have been demonstrated (Clavaguera, et al., 2013), arguing that misfolded tau species in different disease possess different misfolds and therefore different tertiary structure.

Because structural differences between molecules underlie how they interact with and cross cell membranes, such as those in the synapse, tertiary structural differences between misfolded tau strains could account for directional spread biases. Because neuronal signaling and axon structural integrity become progressively degraded over the course of degenerative diseases (Li, et al., 2011), anterograde transynaptic transport likely becomes less frequent and more difficult. This would amount to a selection pressure towards misfolded tau strains that are more readily able to cross the synapse from dendritic bouton to axon terminal, thereby potentially leading to a progressively developing retrograde bias in transcellular and transregional tau pathology spread, across conditions. We posit both above hypotheses as plausible, non-exhaustive, and non-mutually exclusive explanations for our observation of progressively more retrogradely biased tau pathology transmission over time.

### ADL tau datasets show the strongest directional bias at all timepoints

Retrograde bias in tau pathology transmission was observed in two different manners: 1) ADL tau pathology datasets consistently exhibited a retrograde spread bias at all timepoints (Figure 4a), while NADL tau datasets only did so, weakly, at later timepoints, and 2) Across all datasets, tau pathology exhibits a progressive shift towards retrograde propagation, over time (Figure 4b & c). Why do ADL and NADL tau behave differently in these regards? It cannot be due to differences in the speed of proliferation of tau due to ADL status or initial retrograde or anterograde bias status (Figure 4e) as the average βt diffusion constant of best fit in our DNT models, across datasets, a representation of the speed of diffusion, was not significantly different in all cases. Instead, different tauopathic conditions have different tau spread compositions (Kaufman, et al., 2016), hence it is more likely that certain misfolded tau strains innately spread preferentially in anterograde or retrograde directions. Another possibility is that while retrograde bias increases over time, the presence of Aβ causes ADL tau to exhibit this bias earlier and more strongly.

Amyloid could directly alter misfolded tau structure or could indirectly alter tau spread by causing damage to the axon-soma barrier (Bolmont, et.al., 2007; Götz, et.al., 2004; Ittner, et.al., 2010). Of note, however, not all datasets exhibited this progressive shift towards predominant retrograde spread; why some tau pathology is resistant to this shift, and the exact mechanisms that make this shift so prevalent across these datasets should be a source of further inquiry.

### Focal seeding is not necessary to explain regions exhibiting early tau tangle pathology

The presented evidence for retrograde bias in ADL tau transmission opens the possibility that perhaps the early sites of tau pathology (EC, hippocampus, retrosplenial and cingulate cortices) might be better described as retrograde *accumulators* of diffuse pathology rather than focal *initiators* of pathology. We therefore hypothesized that focal seeding would not be necessary to recapitulate the observed patterns of tau deposition, especially in the non-seeded dataset (Hurtado, et al., 2010). Random seeding simulations in Figure 6 show that starting from disparate and random seed locations, subsequent DNT recreated pathology development almost as well as seeding based on baseline pathology in our non-exogenously seeded dataset (Hurtado, et al., 2010). Our results therefore imply that AD pathology can initiate in different loci and still develop with the same spatiotemporal pattern, consistent with a recent human study concluding there may not be specific or consistent initiation loci (Arendt, et.al., 2015). In this view putative “seed regions”, such as the EC or LC or hippocampus, likely initiate tauopathy in a subset, but not all, AD cases, implying proteinopathy need not have a consistent region of initiation. These findings raise an important question: if AD-related tau pathology does not start in a consistent location, what explains the highly stereotyped spatiotemporal development of this proteinopathy?

There is good theoretical support for the hypothesis that regions such as the EC may be accumulators rather than initiators of early tau pathology and that this feature is due to their topographic location in the brain’s network. First, the hippocampal role in the default mode network (DMN) in humans and in mice (Mechling, et.al., 2014) and the EC’s well characterized role as a gate for activity and information flowing from or to the hippocampus (Squire & Zola, 1996) suggest that these areas’ centrality in brain networks underlie their importance in disease pathogenesis. Disruption of hippocampal and cingulate cortex interactions in the DMN during resting state activity is thought to be especially important in AD pathogenesis and proliferation (Greicius, et.al., 2004).

To address this hypothesis we investigated whether the steady-state distribution of global network dynamics of DNT predicted sites of early tauopathy from first principles, without looking at any AD data whatsoever. This turns the seeding question on its head: instead of EC acting as an early site of pathology initiation, we ask whether it could be an early site of pathology accumulation, starting from diffuse or random seeding throughout the brain. Both diffuse whole brain and random seeding followed by retrograde DNT consistently implicated the EC, even at early timepoints, as the region with the highest accumulation to tau pathology (Figure 5b-e). Other regions implicated as early and outsized tauopathy accumulators were early AD regions, including the anterior cingulate, retrosplenial, and piriform cortices; all exhibit early tangle pathology in either AD patients or mouse models (Greicius, et.al., 2004; Ikeda, et.al., 2005; Li, et.al., 2010). Using anterograde DNT we observed the caudoputamen as the earliest and most prominent pathology accumulator (Figure 5b-e).

Since graph diffusion theory predicts that diffusive disease factors will be trapped within spatial patterns that follow Laplacian eigenvectors (Raj, et.al., 2012), we also examined which nodes in the graph have the highest values in *u*_*1*_, the first and most important retrograde Laplacian eigenvector; these were the EC and adjacent piriform cortex. In contrast, the anterograde Laplacian’s first eigenvector showed the highest value in the caudoputamen (Figure 5a). The evidence from *u*_*1*_ of each network together with that from the diffuse and random seeding simulations indicates these regions as the sites likeliest to accumulate enough sub-threshold pathology in each direction to begin the subsequent process of templated corruption (Hardy & Revesz, 2012). Interestingly, the sites implicated as early accumulators of tau pathology in the retrograde network were many of the same areas known to have prominent early AD pathology (Braak & Braak, 1991), while the caudoputamen is thought to exhibit very high early tangle pathology in both tauopathic Parkinsonism and bvFTD (Chiba, et.al., 2012). In all cases, specific seeding in any location, such as the LC or EC, was not necessary to explain regional vulnerability to early tangle pathology. Thus, the seeming SVb of certain brain regions to early tau pathology can be explained purely as a consequence of network topology combined with tau-species or condition specific directional spread biases rather than due to cell-specific, cytoarchitectural or other local, brain region intrinsic and specific properties.

### Conclusions

The present work demonstrates that the diverse spatiotemporal presentation of tauopathic conditions might be explainable simply due to differential directional biases during transsynaptic and transregional spread, an expansion of the MNx hypothesis (Warren, et al., 2013). For example, ADL tau pathology showed a strong retrograde spread bias, while NADL pathology did not, suggesting that AD is differentiable from other tau conditions because of its consistent retrograde bias in pathology spread. Furthermore, we found a general progressively developing retrograde tau spread bias, regardless of tau pathology type or origin. This provides a potential explanation for why AD symptoms, while not appearing early in CBD and FTD cases, eventually appear in these diseases (Burrell, et al., 2016): as these conditions progress, tau spread becomes more dominated by retrograde transmission, akin to AD, and therefore begins to affect many similar regions. Finally, our results do not necessitate the proposed focal pathology seeding due to local regional or specific cell-type properties suggested by the SVb hypothesis (Seeley, et al., 2009). We instead demonstrate that entirely random seeding simulations followed by directionally biased tauopathy spread combined with the mouse brain connectivity network’s wiring properties predict that many of the putative AD tau pathology seed regions are actually those most likely to be pathology accumulators or sinks.

## Acknowledgements

We would like to acknowledge the Allen Institute for Brain Science and in particular their mouse reference and connectivity atlas toolboxes and research teams. Without their exemplary contributions to neuroscience and neuroanatomy, the present work would not be possible.

We would also like to acknowledge the Weill Cornell Neuroscience and Radiology Departments, as well as the Feil Family Brain & Mind Research Institute and the many people who generously support Weill Cornell. AR and CM are supported by grants from the National Institutes of Health (R01NS092802). AR and CM would also like to acknowledge our former and present colleagues from the IDEAL lab.

## Author Contributions

CM and AR designed the models, wrote the analysis code in MatLab, and co-wrote the manuscript. CM performed the lit searches to find the previously published datasets or tables and figures used as the “empirical data” here, performed the analyses to collect results and designed the figures and tables.

## Compliance with ethical standards

### Ethical approval and consent to participate.

All mouse-work datasets in the present work was obtained from datasets originally presented in or with other already published work already declaring that all animal work was done under approved protocols and animal care standards.

## Supplemental Figures

**S. Figure 1.**
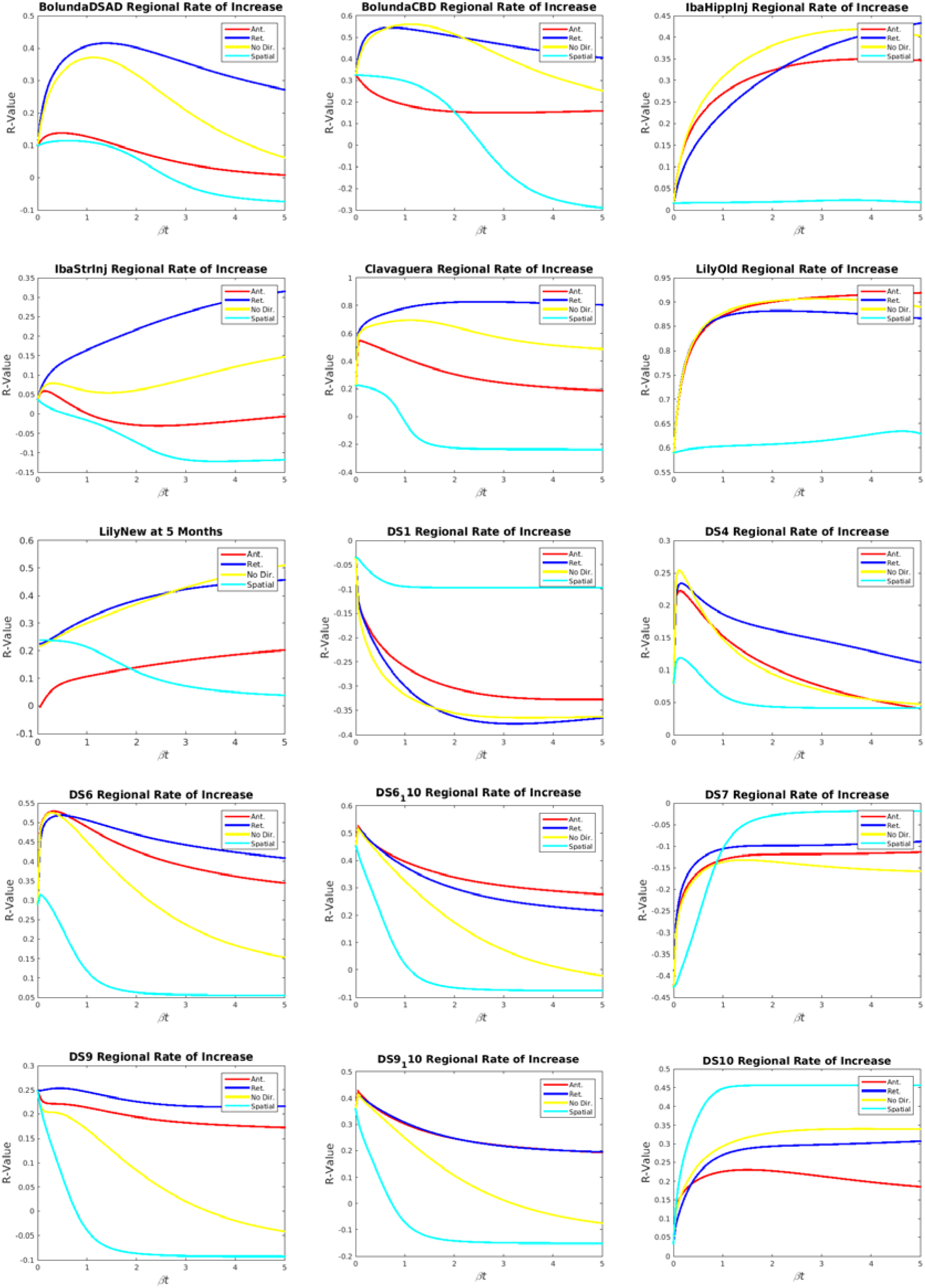
Individual dataset βt diffusion time parameter curves are plotted against r-value of each (anterograde, retrograde, or non-directed) DNT model using that βt value versus empirical pathology regional rate of increase data, with pathology spread initiation starting from reported seedpoint. As in the example illustration in Figure 1b, at βt = 0, this represents the match between the reported seedpoint and the data. The maximum r-value for each line in each graph (anterograde, retrograde, or non-directed) represents the maximum r-value achieved by that model for that dataset. The maximum r-value minus the r-value at βt = 0 represents ΔR, the measurement we use to determine how much predictive value is added by each model, starting from the seecdpoint. Each dataset’s signifier is in the title above the graph. Because the dataset from Hurtado, et al., 2010, did not have any exogenous seeding and the one from Iba, et al., 2015 used a seed region outside the scope of the ABI Mouse Brain Atlas, they are not included here.

**S. Figure 2.**
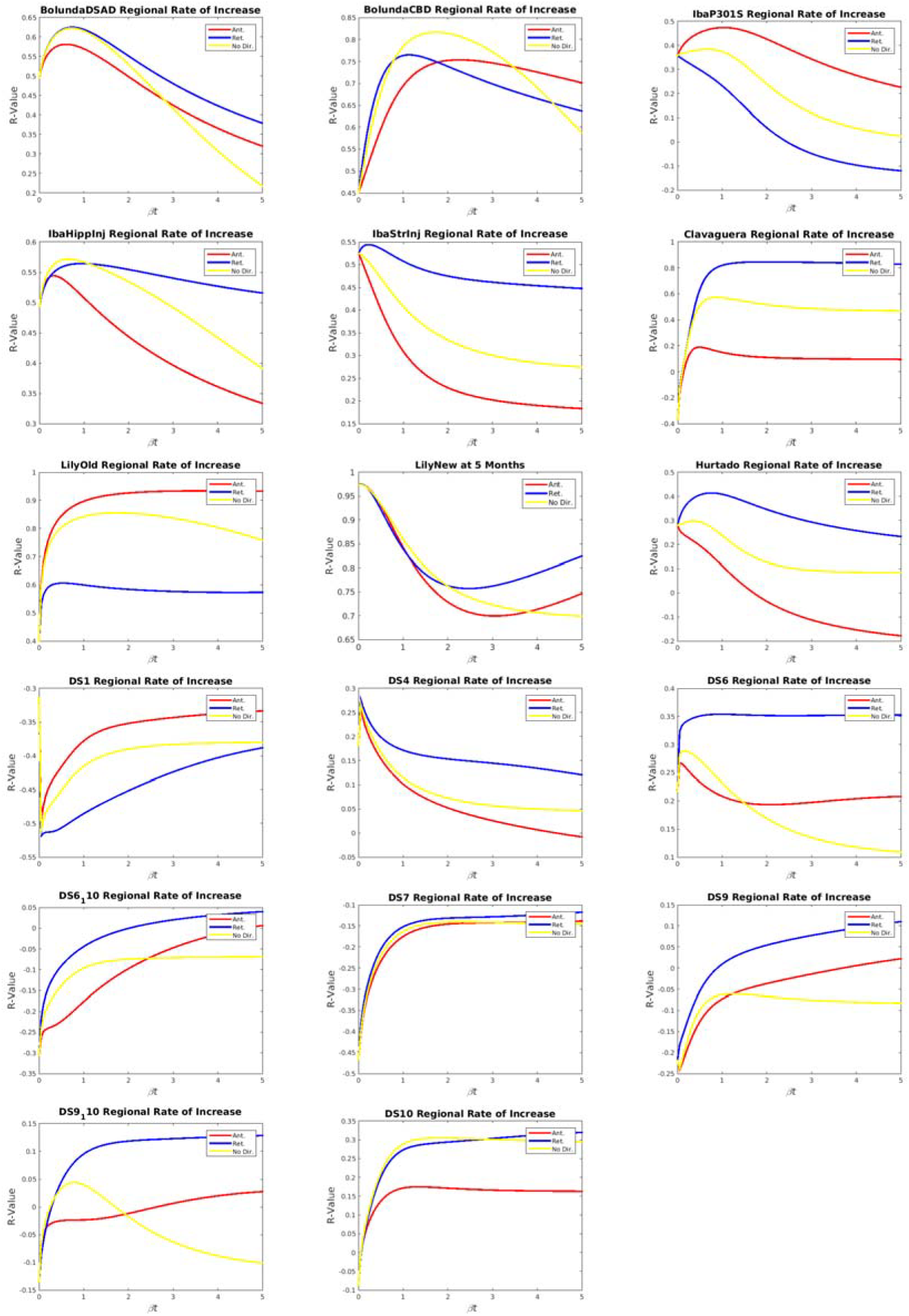
This figure is the same as S. Figure 1, except that pathology is initiated from the per study generated baseline pathology measurement. Because all studies that were longitudinal included a baseline pathology measurement, both the Hurtado, et al., 2010 and Iba, et al., 2015 datasets are included here, unlike in S. Figure 1. Each dataset’s signifier is in the title above the graph.

**S. Figure 3.**
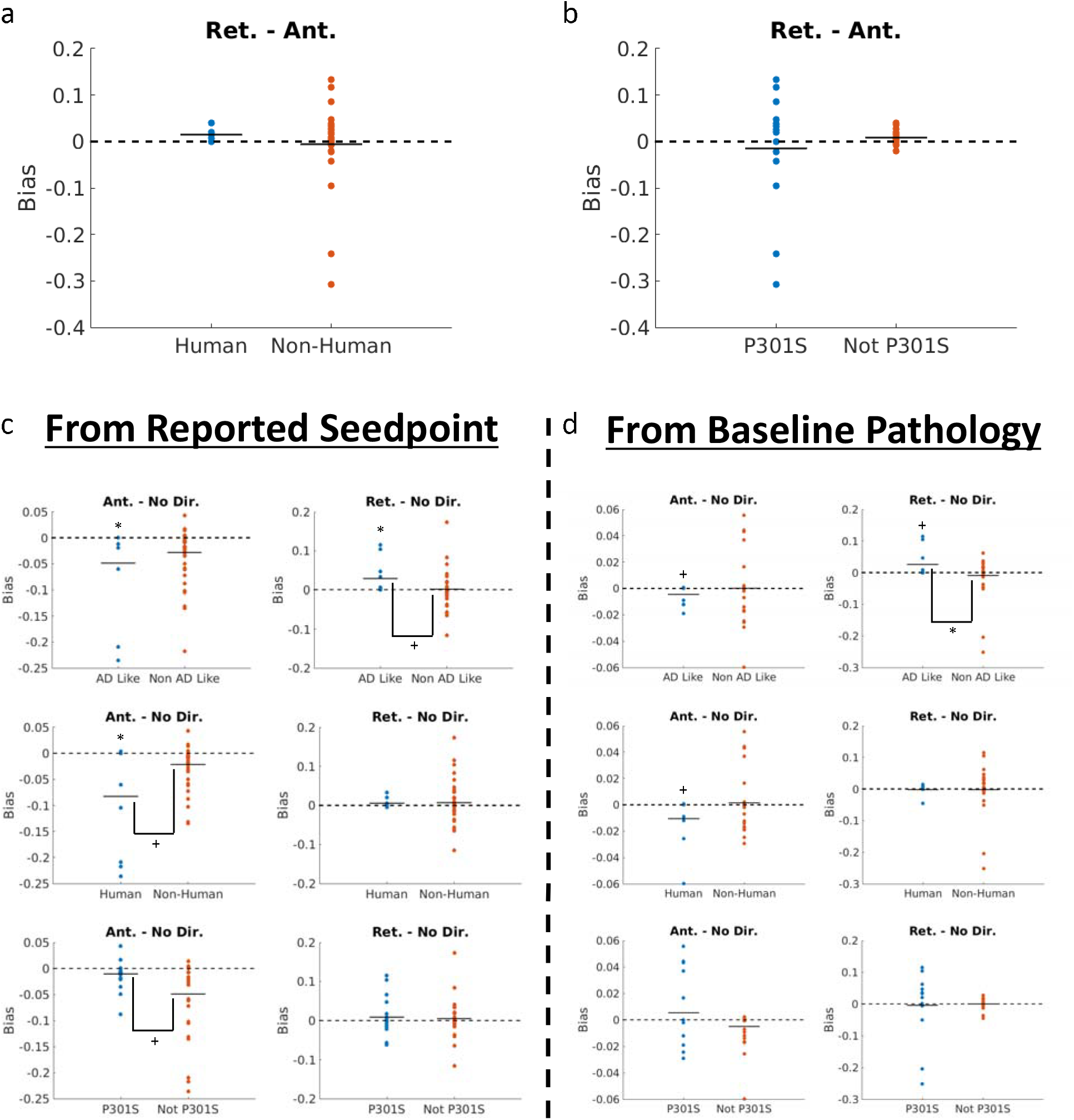
ADL tau shows a consistent retrograde bias in its spread while NADL tau does not, but no other group divisions of the set of studies in the present research elicit any significant results. Both (a) human versus non-human origin of misfolded tau and (b) the aberrant tau being of the P301S versus some other variety demonstrate no effect in terms of spread bias in any direction. Modeling tau pathology spread from both reported seedpoint (c) and from the baseline regional pathology measurement (d), across studies and timepoints, demonstrates that ADL tau in mouse models exhibits a consistent and significant (or trending towards) retrograde bias relative to nondirectional spread, but no other groupings of studies yield significant results. The asterisks and crosses above a distribution indicate that the distribution is significantly different from 0 in a one-sample t-test and the asterisks and crosses below the brackets hanging downwards between each pair of distributions indicates the distributions are significantly different from each other in a two-sample t-test.

**S. Figure 4.**
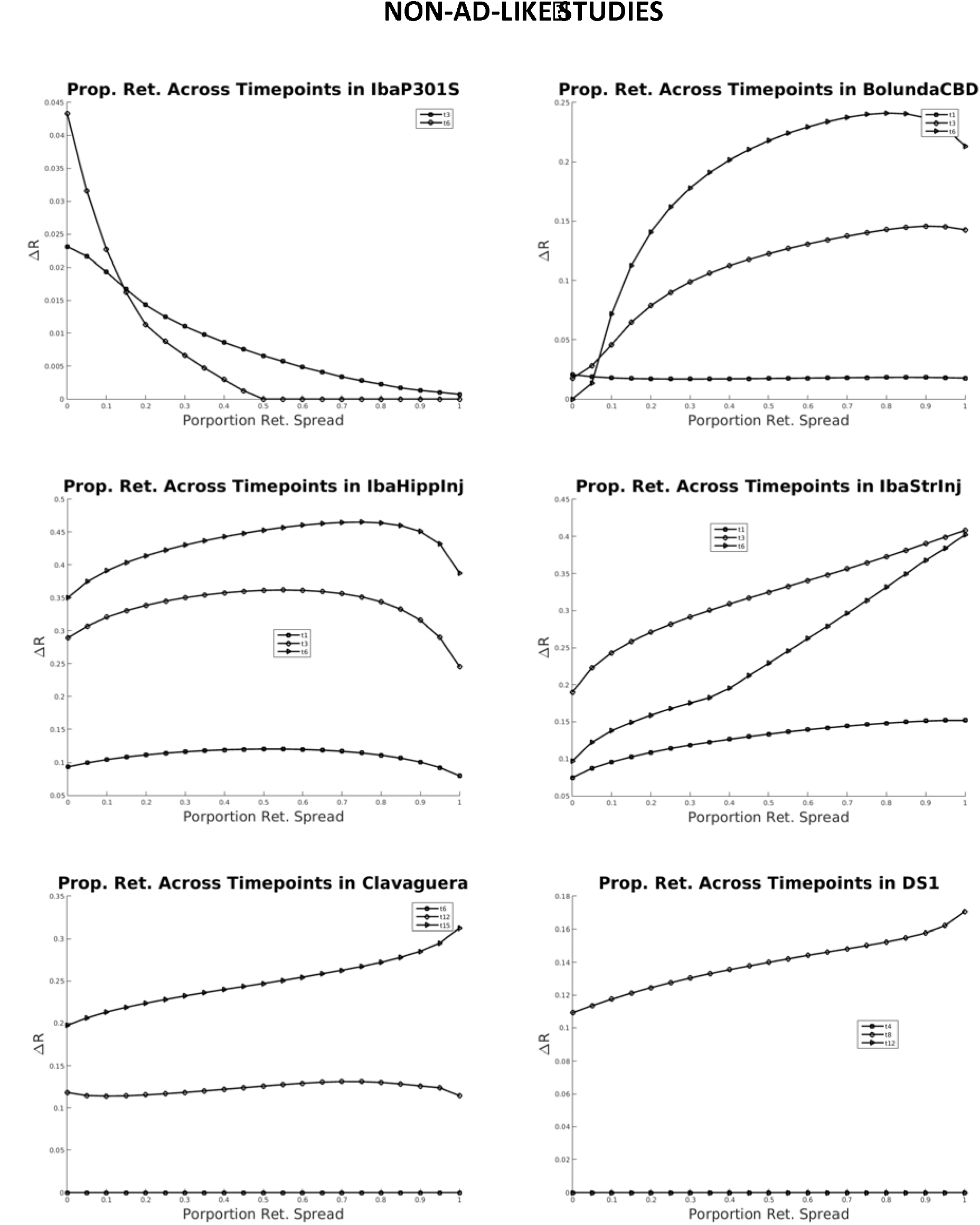
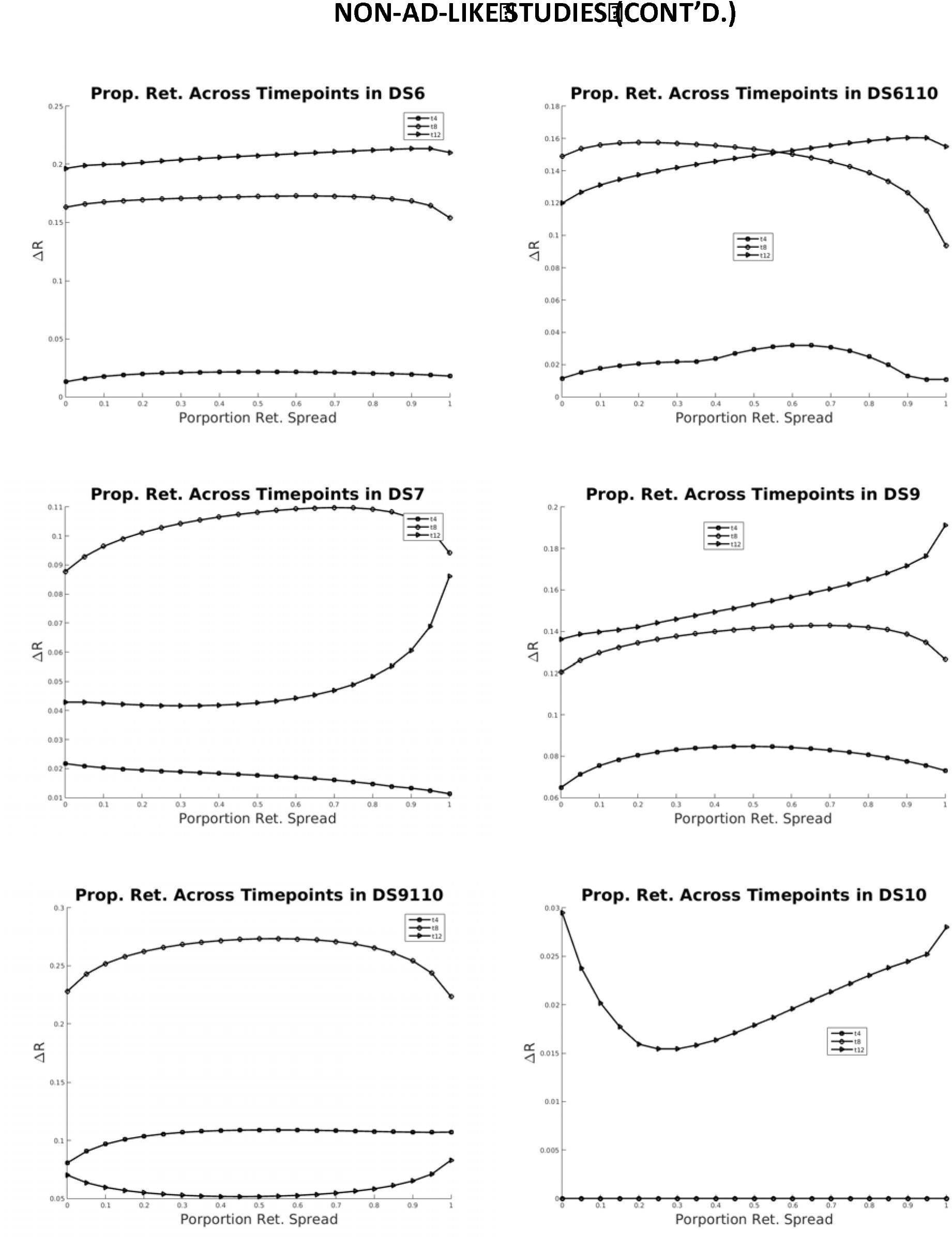
Here we exhibit the proportionate DNT model, across all timepoints in NADL datasets, with the dataset in question as the title of each plot and each curve in each plot representing the different measured timepoints in each study. Each curve is the proportion of retrograde spread in the DNT model used here against the R-value with the timepoint and dataset in question. The higher the x-axis value, the higher the proportion of retrograde spread, and the higher the y-axis value, the higher the R-value. Note: this figure spans two pages as we have too many NADL datasets to plot onto one page.

**S. Figure 5.**
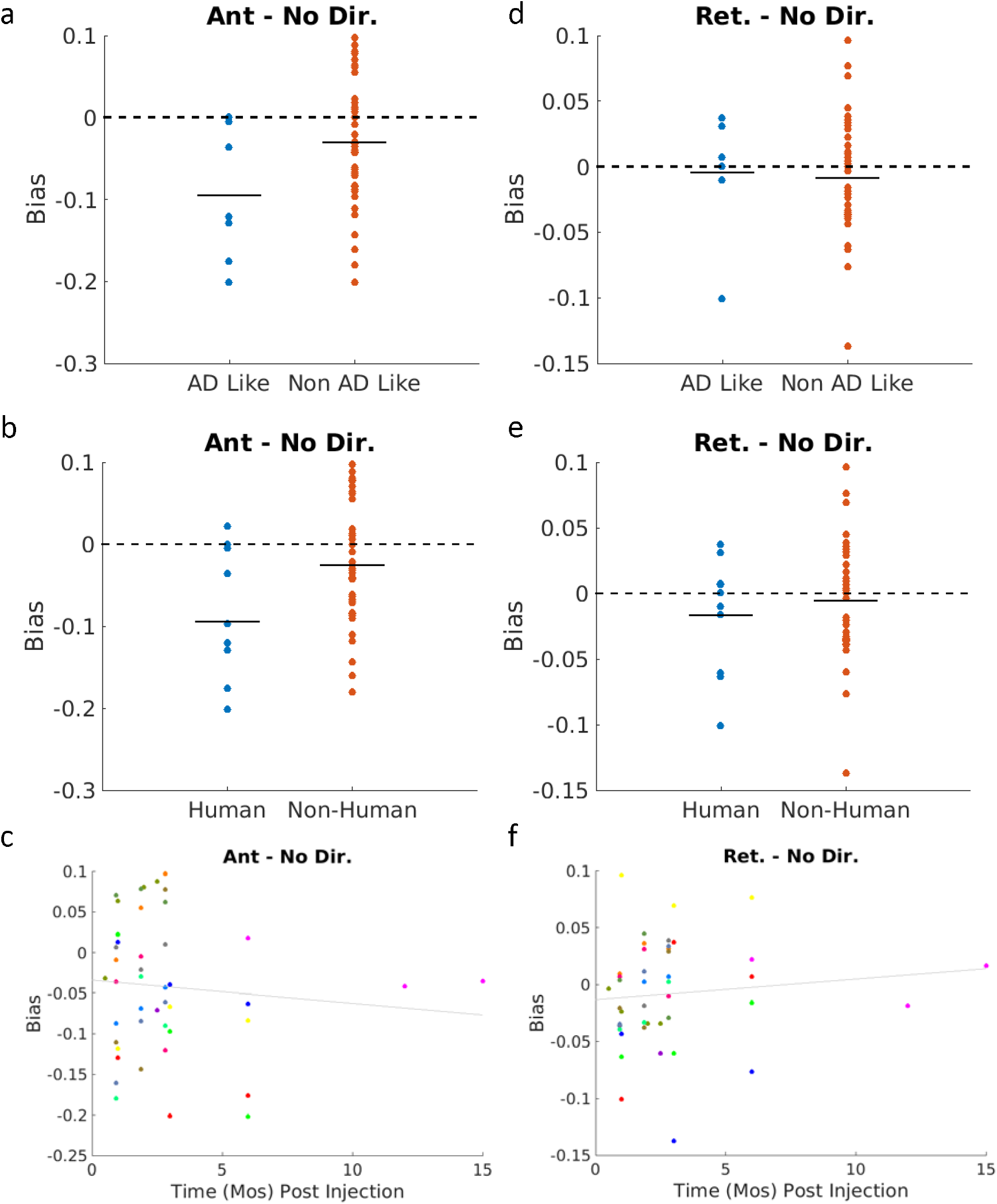
Retrograde and Non-Directional, but not anterograde, connectivity with reported seed regions are predictive of empirical tau pathology patterns. (a) ADL and (b) human origin tau utilizing datasets show a bias away from anterograde connectivity being predictive of empirical tau pathology patterns. (c) Anterograde connectivity with reported tau pathology seed regions becomes progressively worse at recapitulating empirical tau pathology. (d) ADL tau datasets show a slight shift towards a retrograde connectivity with reported seed region being more predictive of empirical tau pathology, a pattern akin to that observed using retrograde DNT, albeit not as strong. (e) Human origin and non-human origin tau show no directional bias in how predictive connectivity with the seed region was of empirical tau pathology patterns. (f) Retrograde connectivity with the seed region becomes progressively better at recapitulating empirical tau pathology progression over time, akin with the results using retrograde DNT. In both (c) and (f) each color represents a study, so studies can be tracked over time in these panels.

**S. Figure 6.**
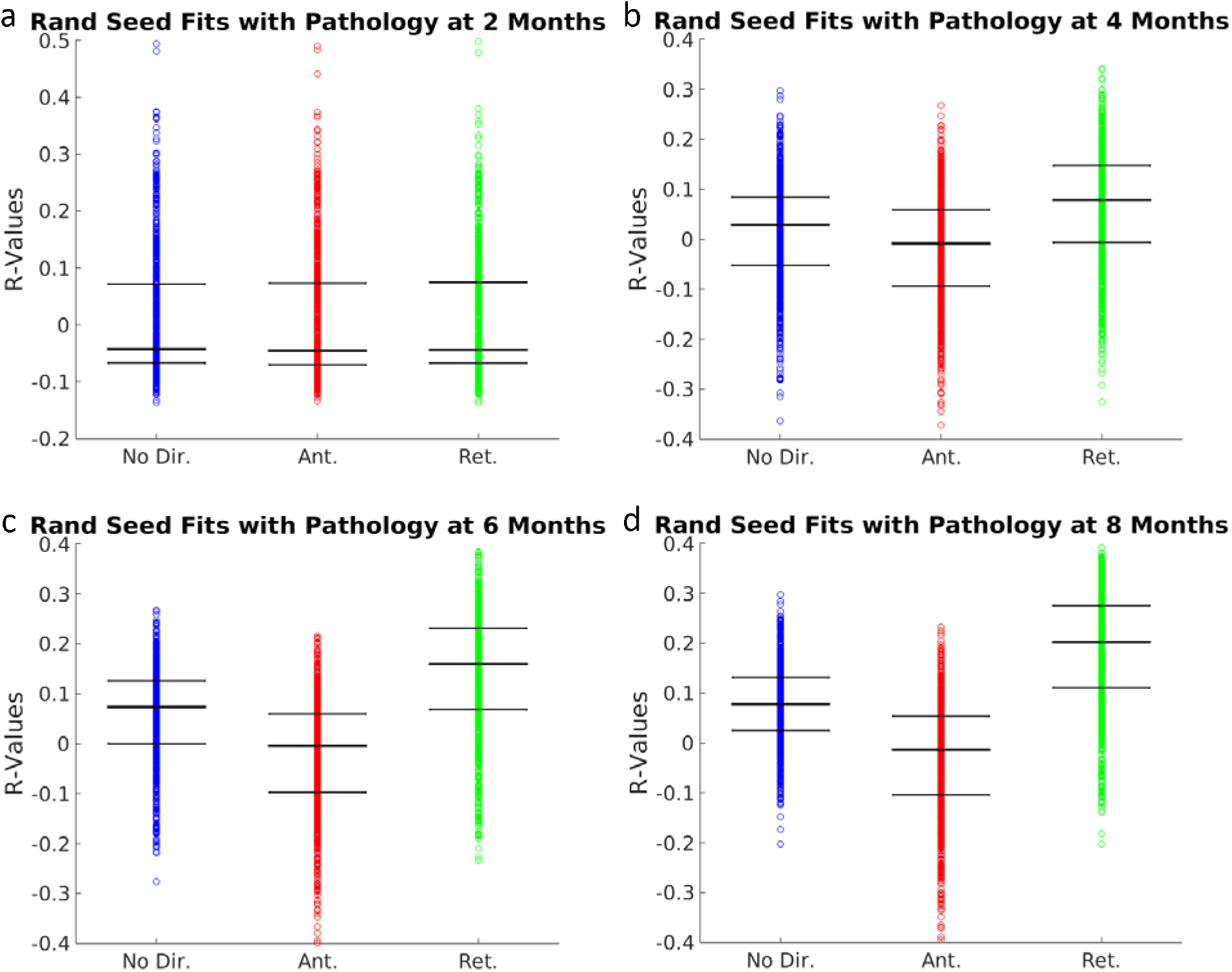
Random seeding simulations followed by retrograde DNT, but not anterograde or nondirectional DNT, produce higher R-values with empirical data from the ADL nonseeded Hurtado, et al., 2010 dataset. This pattern was not observed at (a) 2 months post birth of the APPswe x P301S mice, but was evident by (b) 4 months post birth, and was maintained at both (c) 6 months and (d) 8 months post birth of the transgenic animals.

